# AI-Powered Acoustic Surveillance for Early Detection of Calf Respiratory Disease

**DOI:** 10.64898/2026.01.20.700576

**Authors:** Núria Mach, Ignasi Nou-Plana, Marie Corbin, Mariette Ducatez, Gilles Meyer, Rosa Maria Alsina-Pagès, Antonio Velarde

**Affiliations:** Univ Toulouse, ENVT, INRAE, IHAP, Toulouse, France; Human Environment Research (HER), La Salle - Universitat Ramon Llull; IRTA, Animal Welfare, Monells, Catalonia, Spain

**Keywords:** AI-powered acoustic monitoring, bovine respiratory disease complex, early warning, cough, precision livestock farming

## Abstract

Effective management of Bovine Respiratory Disease Complex (BRDC) requires timely, non-invasive diagnostic tools to protect calf health and welfare. Among early clinical signs, coughing stands out as both frequent and informative. To explore its potential for early BRDC detection, we deployed an artificial intelligence (AI)-driven acoustic monitoring system that recorded over 2,730 hours of audio during a 30-day period. Four experimental pens, each housing seven calves and stratified by infection status and antibiotic treatment, were equipped with a dedicated microphone to enable targeted acoustic surveillance. This configuration enabled pen-specific detection of cough events, which were subsequently classified using an AI HuBERT-based model trained on 1,045 labelled clips. The classifier achieved 92% accuracy. Temporal patterns in cough frequency aligned with infection dynamics, treatment responses, and circadian patterns. Notably, AI-detected coughs consistently preceded clinical scores by 1–2 days, confirming the system’s sensitivity to early respiratory disorders. These findings support the use of acoustic surveillance as a valid, scalable, and autonomous tool for continuous monitoring and early warning of respiratory diseases in calves.

**Implications:** This study demonstrates that AI-powered acoustic monitoring enables real-time, non-invasive detection of coughs in calves for early warning of respiratory diseases, outperforming traditional veterinarian clinical scoring by 1–2 days. Its high accuracy and sensitivity to respiratory infection dynamics and treatment effects position it as a scalable tool for precision livestock farming.

## Introduction

Bovine Respiratory Disease Complex (BRDC) is one of the most significant health and welfare challenges in beef cattle production worldwide (Gaudino et al., 2022; Taylor et al., 2010; Timsit et al., 2016), and is also a major driver of antimicrobial use. This multifactorial syndrome arises from the interaction of environmental stressors, host-associated (holobiont) factors, and a diverse range of respiratory pathogens (Arcangioli et al., 2008; Gaudino et al., 2023; Griffin et al., 2010a; Holman et al., 2015; Klima et al., 2019; Salem et al., 2019a, 2020). Young calves are particularly susceptible, as BRDC can impair immune development, induce pain and distress, and substantially increase the risk of long-term morbidity and mortality (Cummings et al., 2022). This is particularly relevant during transfer to fattening units, where calves from different origins are mixed, exposed to novel pathogens, and subjected to additional stress.

BRDC early detection is essential—not only to reduce economic losses, but, more importantly, to ensure timely, targeted interventions that increase treatment success, prevent suffering, and thereby protect animal health and welfare (Nielsen et al., 2023). BRDC is clinically characterized by signs such as coughing, sneezing, nasal discharge, and tachypnoea (Carpentier et al., 2018; Gaudino, Nagamine, et al., 2022), with coughing being a particularly informative symptom for early detection (Carpentier et al., 2018). According to EFSA (2025), the sensitivity and specificity of coughing as an indicator of respiratory disorders are high (Nielsen et al., 2025). Coughing results from a forceful expulsion of air from the lungs triggered by a sudden contraction of the diaphragm and intercostal muscles in response to irritation of the lower respiratory tract (Nielsen et al., 2025). Variations in cough type may be associated with different respiratory conditions, affecting either the upper or lower tract, and may occur with or without fluid accumulation. The Welfare Quality protocol (2023) for fattening cattle recommends quantifying cough frequency by continuous observations of pens or, for very large pen sizes, by monitoring subgroups of no more than 25 animals (Welfare Quality Network, 2023). However, the expansion of large-scale farming systems, combined with increasing labour constraints, renders widespread adoption of manual coughing assessments difficult to implement in practice. Furthermore, traditional BRDC diagnostic methods, which rely on pathogen-oriented sampling during farm visits (Werid et al., 2024), are costly, invasive, and potentially stressful, and are often performed with a delay, which limits their effectiveness in detecting early disease onset (Zhang et al., 2022).

These limitations have accelerated the development and adoption of Precision Livestock Farming (PLF) technologies, which use automated data collection to support on-farm decision-making and are becoming increasingly important for early disease detection in intensive livestock production systems (Norton and Berckmans, 2018). Cough monitoring technologies have been developed to support on-farm early detection of respiratory disease in livestock (Berckmans et al., 2015). In pigs, algorithms using historical data and room-specific variations generated a respiratory distress index that correlates moderately (r=0.5) with manually recorded coughing frequencies (Pessoa et al., 2021). The same study further showed that this index, measured at the end of the finishing period (21–24 weeks of age), correlated strongly (r > 0.7) with the prevalence of lung lesions and pneumonia assessed at slaughter. In cattle, Vandermeulen et al. (2016) reported high specificity (99.2%) and precision (87.5%) in calf cough detection, while Carpentier et al. (2018) demonstrated field applicability with precision exceeding 80% in several compartments and a clear correlation between cough frequency and BRDC diagnosis. Complementary approaches using collar-mounted sensors also achieved strong performance metrics, including 88.9% accuracy (Kim et al., 2024), reinforcing the viability of acoustic and behavioral monitoring for early disease surveillance.

While these results support the potential of digital farming sensors for cough as an automated tool for continuous detection of respiratory disorders in calves, further optimization is required to improve specificity and scalability in precision livestock systems. In this context, the COVID-19 pandemic served as a powerful accelerator for acoustic cutting-edge surveillance technologies, demonstrating how AI-powered cough detection systems could be deployed at scale for early diagnosis and outbreak monitoring (Askari Nasab et al., 2023; Ayappan & Anila, 2025; Feng et al., 2024; Pentakota et al., 2023). These developments—encompassing refined cough signal processing, machine learning and deep learning algorithms, and remote health monitoring infrastructures—provide a robust translational framework for veterinary medicine. Therefore, building on these advances in human medicine, our study aims to adapt and validate acoustic surveillance methodologies for use in young ruminants under controlled experimental conditions. Calves were continuously monitored from arrival, with respiratory pathogens inoculated at a defined time point to control disease onset. Cough events captured via strategically placed microphones were systematically compared with cough assessments performed by veterinarians. This design enables precise temporal alignment between acoustic signals and true disease onset, thereby strengthening the reliability and diagnostic value of cough-based monitoring systems.

## Material and Methods

### Animal handling and groups at the experimental facility

Following birth at INRAE-Herbipole, 28 healthy and unweaned calves (30.7□±□13.3 days old) were transported to the national veterinary school in Toulouse (ENVT, Toulouse, France). Animals were assigned to four groups: (i) infected without antibiotics (*Inf_NonATB*), (ii) infected with antibiotics (*Inf_ATB*), (iii) uninfected with antibiotics (*NonInf_ATB*), and (iv) uninfected without antibiotics (*NonInf_NonATB*). Groups were balanced by sex, age, breed, and body weight.

Antibiotic treatment with tilmicosin (12.5□mg/kg, twice daily, Tilmovet 250 mg/mL, Huvepharma, Antwerp, Belgium) was administered from day -5 to -3 of the experiment. On day 0, calves in the infected groups were co-inoculated with three respiratory pathogens: Influenza D virus (IDV, strain D/bovine/France/5920/2014), Bovine Coronavirus (BCoV, strain ENVT-66), and *Mycoplasma bovis* (strain RM16) (Lion et al., 2021). Inoculation was performed via intranasal nebulization (8–12□min) using a foal-adapted mask and compressor system, delivering 10□ TCID_50_ of IDV and BCoV, and 10^1^□ CFU of *M. bovis* per calf. Prior to inoculation, all calves tested negative for major respiratory pathogens and antibodies.

The study took place in a biosafety level 1 barn (35□×□11.3□×□5□m) with natural ventilation and four pens (4.9□×□14□m; 10.5□m^2^/calf), arranged in two rows separated by a 1.5□m corridor (Fig.□1AB). Each pen housed seven calves and featured open sides, solid rear walls, and a galvanized mono-pitch roof. Infected and uninfected groups were separated by a transparent polyethylene barrier, allowing visual but not tactile contact. Pens operated independently under a “*marche en avant*” protocol, with no shared equipment or personnel crossover. Bedding (pine shavings) was spot-cleaned daily and replaced every two days.

**Figure 1.**
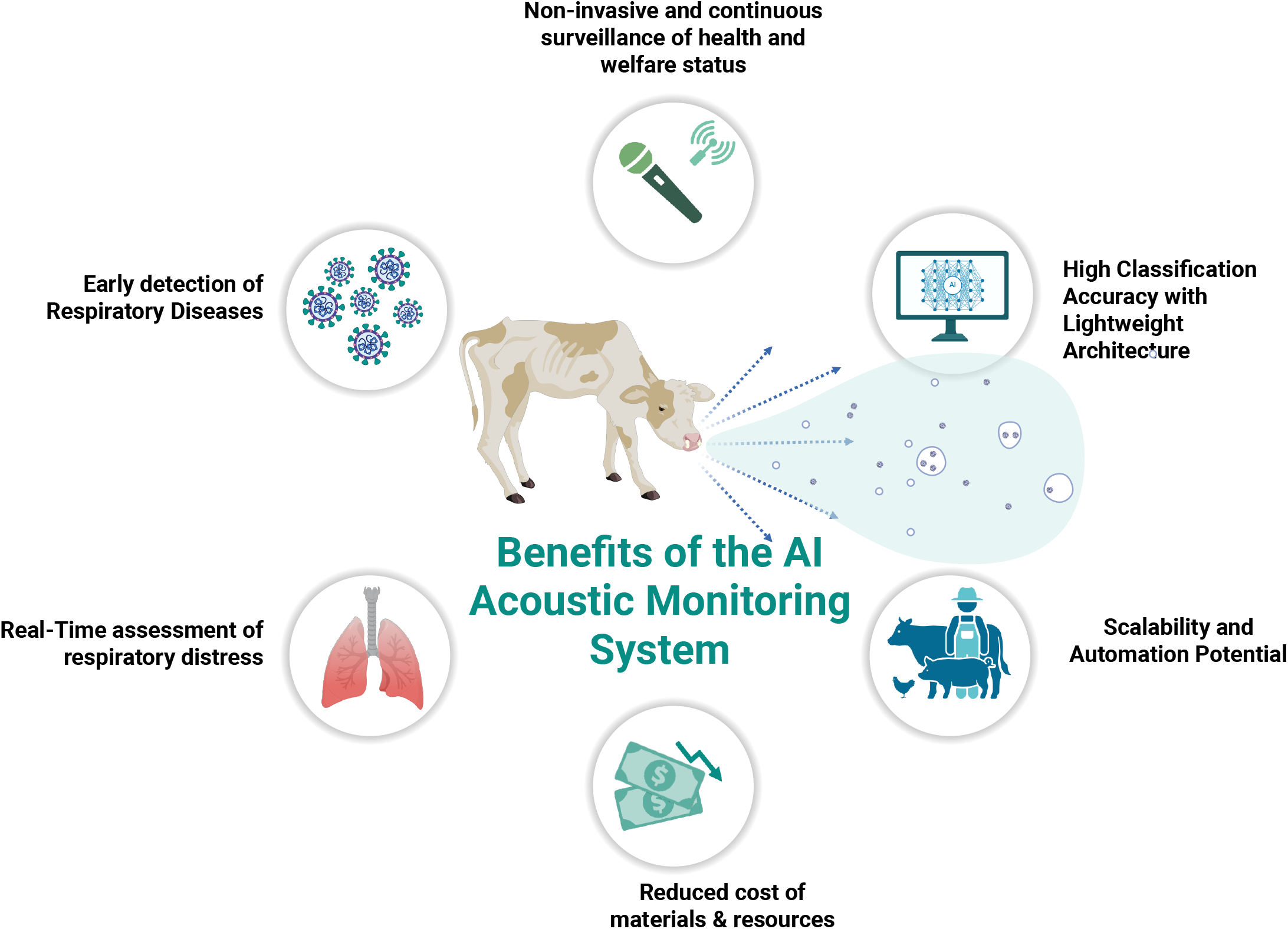
Experimental Setup and Clinical Cough Monitoring in Calves. (A–B) show pen layout and group allocation: 28 calves were divided into four groups (*Inf_ATB, Inf_NonATB, NonInf_ATB, NonInf_NonATB*), each housed in a microphone-equipped pen for continuous cough monitoring; (C) summarizes pathogen exposure (IDV, BCoV, *Mycoplasma bovis*) and antibiotic protocol; (D) Project timeline: calves arrived 12 days before infection (Day 0), with monitoring continuing until Day 19 via twice-daily clinical scoring and uninterrupted audio recording; (E) Smoothed mean cough scores by day post-infection (Day 0), shown for each experimental group; (F) Daily pairwise comparisons highlight significant group differences. Tile colour intensity reflects –log10(*p*-value), with darker reds indicating stronger significance; asterisks mark *p*□<□0.05. Pale or grey tiles denote nonsignificant or missing data.

Milk replacer was offered twice daily (07:00 and 17:00), with a 30-minute feeding window. The quantity of milk refused was recorded without force-feeding. The milk replacer (Univor, Spécial Élevage, France) was adjusted weekly to support a target growth rate of 0.8□kg/day. Initial volumes ranged from 3–4.5□L per meal and increased to 4.7–6.7□L based on body weight. Calves had *ad libitum* access to fresh water via automatic drinkers. Neither concentrate nor hay was provided during the study.

### Clinical cough assessment by veterinarians

From arrival (Day -11) to 19 days post-infection, calves were individually examined twice daily for clinical signs including general condition, appetite, rectal temperature, nasal discharge, respiratory effort, cough, and lung sounds. To minimize the risk of cross-contamination, two veterinarians were assigned exclusively to the two infected pens and two others to the two non-infected pens, with no interchange between groups throughout the study. For this analysis, only cough presence or absence was retained and further examined. During each clinical visit, coughing was assessed by experienced veterinarians using a standardized respiratory examination protocol. Calves were first observed quietly at rest to identify spontaneous coughing events. This was followed by thoracic auscultation, during which both lung fields and the trachea were examined for abnormal respiratory sounds, including harsh bronchial sounds, tracheal rattles, wheezes, or crackles. Each calf was monitored for a defined period to ensure consistent classification. Animals showing spontaneous, or presenting abnormal tracheal or lung sounds on auscultation, were classified as “cough”, whereas those without any of these signs were classified as “no cough”. Each calf was monitored twice daily by two veterinarians, once in the morning and once in the afternoon, to ensure consistent and reliable classification across time. This multiple annotation approach was implemented to independently verify potential bias and ensure that all labelled events corresponded to true coughs.

### Environmental data

Automated sensors (VelociCalc® 9565-A, TSI, USA) recorded temperature, humidity, heat index, and dew point every 10 minutes in each pen.

Dew point and heat index thresholds were predefined to assess thermal stress: <10□°C (“Dry & Crisp”) and >21□°C (“Tropical & Oppressive”) for dew point; >27□°C and >32□°C for heat index (“Caution” and “Extreme Caution”).

### Cough dataset collection using audio

Sound was recorded with dynamic supercardioid microphones (Behringer XM1800S, Willich, Germany) fitted with foam windscreens to protect against wind and dust and to allow quick hygienic replacement. Microphones were hung from the ceiling, about one meter above the calves’ height. The placement of microphones in each pen aimed to maximize their separation while maintaining sufficient distance from walls to reduce unwanted sound reflections. A Raspberry Pi 4 Model B (Cambridge, UK) enabled real-time remote monitoring of input levels. Microphone signals passed through a Focusrite preamp/interface (Scarlett 2i2 USB Audio Interface, Wycombe, UK) and were captured on a ZOOM H4n Pro Handy Recorder (Zoom corporation, Tokyo, Japan) following protocols established in previous related studies (Alsina-Pagès et al., 2021; Miranda et al., 2024; Vidana-Vila et al., 2023).

### Preprocessing of the audio-based Cough Pipeline

From audio recordings, a dataset for cough/non-cough classification was built. Eighty manually labelled clips were first used to train an initial classifier. Model predictions on new segments were then used to accelerate annotation: high-confidence positives and negatives were flagged for human verification. This semi-supervised loop—training, inference, assisted labelling, and dataset expansion—was repeated multiple times, increasing both the quantity and diversity of labelled data and steadily improving model performance and annotation efficiency.

All audio files were pre-processed using the Torchaudio library (Hwang et al., 2023). Recordings at 48 kHz were resampled to 16 kHz to match the pre-trained lightweight Hidden-Unit BERT (HuBERT) front end (Hsu et al., 2021). To improve robustness under field conditions, notch filtering was optionally applied to attenuate power-line interference at 50 Hz and low-order harmonics (e.g., 100, 150 Hz). Band-pass filters (Q = 3) were implemented using forward–backward (zero-phase) IIR filtering to prevent phase distortion. When augmentation was enabled, both the original and the filtered versions of each clip were retained for training, and the train/test split was performed at the source-file (or pen/day) level to prevent leakage between sets.

### Audio-based Cough Feature Extraction and lightweight HuBERT-Based Architecture

The pretrained Facebook HuBERT-base-ls960 model was employed, a self-supervised speech-representation architecture that generates 768-dimensional embeddings directly from raw audio waveforms. HuBERT is particularly well-suited for cough event detection because it learns acoustic structure, enabling it to capture subtle temporal and spectral patterns characteristic of respiratory sounds. Unlike traditional handcrafted features, HuBERT embeddings encode rich, hierarchical representations of both short-term phonetic cues and longer-range acoustic dynamics, which enhances the model’s ability to discriminate coughs from background noise, vocalisations, and other barn-level acoustic events. Embeddings were obtained by averaging the final encoder layer over time and fed them to a lightweight classifier. Freezing HuBERT reduced computational cost and mitigated overfitting, given the limited volume of labelled data.

### Model Architecture of the audio-based cough classification and training strategy

The classifier operated directly on the 768-dimensional HuBERT embeddings. To minimize overfitting, we used a single fully connected linear layer mapping to two classes (cough vs. non-cough). A softmax function converted outputs to class probabilities; only the classifier’s parameters were updated during training. This minimalist design enabled fast iteration within the semi-supervised workflow, where model predictions assisted progressive data labelling and dataset growth. The training was carried out with cross-entropy loss and the Adam optimizer.

### Inference of cough events from audio-based data

During inference, WAV files are segmented into non-overlapping 1-s windows. Segments were converted to tensors with Torchaudio, afterwards resampled to 16 kHz, then normalized with the Wav2Vec2FeatureExtractor, and passed through the frozen HuBERT encoder (eval mode) to generate 768-dimensional embeddings. The trained linear classifier outputs class probabilities: segments were labelled “cough” when the predicted classification was *cough* and the associated probability exceeded a predefined threshold (default 0.50). Predictions were saved as timestamped text files for each input audio. The pipeline supported batch processing of user-specified folders and runs on CPU, Apple Silicon (MPS), and CUDA-enabled GPUs.

### Performance Assessment of the HuBERT Cough Detection Model from audio-based data

The final model was trained and evaluated on a fully labelled set of 1,045 clips (737 non-cough; 308 cough). All clips were manually inspected to ensure label quality. Standard metrics—precision, recall, F1-score, accuracy, and macro/weighted averages—computed per class and overall were reported to ensure a balanced assessment of performance.

Precision, defined as TP/(TP+FP), is the proportion of predicted positives that are true and was computed separately for the cough and non-cough classes. Higher precision indicates fewer false positives. Recall (sensitivity), defined as TP/(TP+FN), measures the proportion of actual cough events correctly identified. Higher recall corresponds to detecting a larger share of true coughs. These metrics were computed for both classes to ensure balanced evaluation across the dataset. The F1-score, which harmonizes precision and recall, was used to summarize overall classification performance.

### Acoustic alarm threshold estimation and visualization from audio-based data

To detect anomalous increases in cough activity, an alarm threshold was defined based on pre-infection acoustic data. For each microphone, the baseline cough activity was calculated using event counts recorded during the five days preceding the infection date (Day 0). Specifically, the baseline was computed as the mean number of daily cough events plus two standard deviations (*μ* + 2*σ*), where *μ* is the mean baseline cough count (Days –5 to 0), and *σ* is the corresponding standard deviation. Daily cough event counts were then compared against this threshold. Days exceeding the threshold were flagged as alarm-positive, indicating potential early respiratory distress. These alarm flags were visualized using time-series plots, with the *x*-axis representing experimental days relative to infection, and the *y*-axis showing event counts. A red dashed line denoted the alarm threshold, while alarm-positive days were highlighted with red markers and annotated as “Alarm.” All plots used a fixed *y*-axis scale to facilitate cross-sensor comparison. This approach enabled biologically relevant and interpretable detection of acoustic anomalies, supporting early intervention strategies in precision livestock monitoring.

### Statistical Analysis: Temporal Dynamics of Cough Frequency Across Treatment Groups

Temporal changes in cough frequency were assessed using two complementary datasets: clinical scores from veterinarian evaluations and event-level predictions from the automated audio-based pipeline.

#### Veterinarian-Assessed Cough Scores

Statistical analyses were performed in R to evaluate group-wise differences over time. A linear mixed-effects model was applied to the full dataset using the *lmer()* function, with cough score as the dependent variable, treatment group as a fixed effect, and animal ID as a random intercept to account for repeated measures. To explore daily contrasts, the dataset was grouped by day, and separate linear models were fitted for each subset. Post-hoc pairwise comparisons were conducted using the emmeans package (v1.10.0) and Tukey’s adjustment for multiple comparisons.

#### Audio-Inferred Cough Events

Each audio-detected event was timestamped and associated with a continuous probability score. For statistical modelling, each row represented a discrete event, and daily counts per sensor were aggregated using the *aggregate()* function in R. The resulting dataset included total event counts (Event_Count) per date and group. To assess temporal and group-wise differences in cough detection, a generalized linear model (GLM) with a Poisson distribution and log link function was fitted. Predictor variables included treatment group and recording day, with the reference group set to *NonInf_NonATB*. Statistical significance was evaluated using *z*-statistics and associated *p*-values, with a threshold of *p* < 0.05.

### Comparative Analysis of Cough Detection: Audio-based vs. Veterinarian Observations

Longitudinal cough data collected via time-stamped microphone recordings were compared against veterinarian-assigned clinical scores. To enable direct comparison, both datasets were aggregated at the group × date level.

Because the two metrics differ in scale, veterinarian cough scores were rescaled to align with audio-based event frequency. This transformation enabled graphical overlay and unified statistical modelling of both measures.

To explore the relationship between scaled veterinarian cough scores and cough audio-based event frequency, a generalized additive model (GAM) was fitted using restricted maximum likelihood (REML) smoothing:

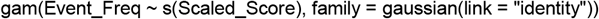

This approach allowed for flexible, non-linear modelling of the association between cough veterinarian scores and audio-derived metrics. Model performance was evaluated using adjusted R^2^, deviance explained, and the significance of the smooth term.

## Results

### Temporal dynamics of Respiratory Cough frequency Across Treatment Groups as assessed by the veterinarians

The progression of cough scores following infection and antibiotic administration was evaluated by veterinarians from Day –11 to Day +19 post-infection across the four experimental groups. Smoothed trajectories of mean cough scores revealed distinct temporal patterns linked to infection status and treatment (Fig. 1EF; lmer, *Group*, F_1,1672_= 62.16, *p* < 0.001; *Day*, F_1_,_1672_= 62.54, *p* < 0.001, and their interaction *Group × Day*, F_3_,_115_ = 24.58, *p* < 0.001). Respiratory coughs emerged around Day +5 in experimentally infected calves without antibiotic treatment (*Inf_NonATB*), marking the earliest onset among all groups. Cough scores continued to rise and peaked at Day +11, reflecting the infection’s most pronounced clinical expression. In contrast, infected calves treated with antibiotics (*Inf_ATB*) demonstrated a consistent therapeutic response. From Day 9 onward, their cough scores were significantly lower than those of untreated infected calves (e.g., Day 9: estimate = –0.57, *p*□=□0.0366), confirming the efficacy of antimicrobial intervention (Fig. 1F).

Unexpectedly, the *NonInf_NonATB* group—intended as untreated, uninfected controls—also displayed notable episodes of coughing, particularly in the mid-phase of the study, albeit less pronounced than in the *Inf_NonATB* group. These cough events likely reflect cross-contamination or natural transmission from experimentally infected animals, despite physical separation and biosafety precautions. Non-infected calves receiving antibiotics (*NonInf_ATB*) maintained consistently low respiratory scores throughout the observation period.

Beyond infection-related dynamics, most groups exhibited an early surge in cough events during the adaptation period. This peak likely reflected group stress—mixing unfamiliar animals by weight, sex, and breed increases activity (e.g., agonistic interactions), vocalization, and respiratory effort (Lyu et al., 2023). Additionally, initial exposure to fresh pine shavings may have caused airway irritation from dust, previously linked to coughing in housed calves (Donlon et al., 2023). Importantly, this transient increase in cough activity was not associated with changes in ambient temperature or humidity, which remained stable throughout the study period (Fig. S1). Nor was it linked to dietary transitions, as animals were not exposed to hay or concentrate at any point during the trial and remained on the same milk replacer (Fig. S1).

### High-Performance Acoustic Cough Classification Using Lightweight HuBERT-Based Architecture

Over the study, the four sensors recorded 2,730.7 hours of audio across four calf pens and ∼30 days (Fig. 2A). Of the 482 files generated with an average duration of 5.7 hours per file, 89.0% were successfully processed. Only 53 were discarded due to corruption or absence of detectable events.

**Figure 2.**
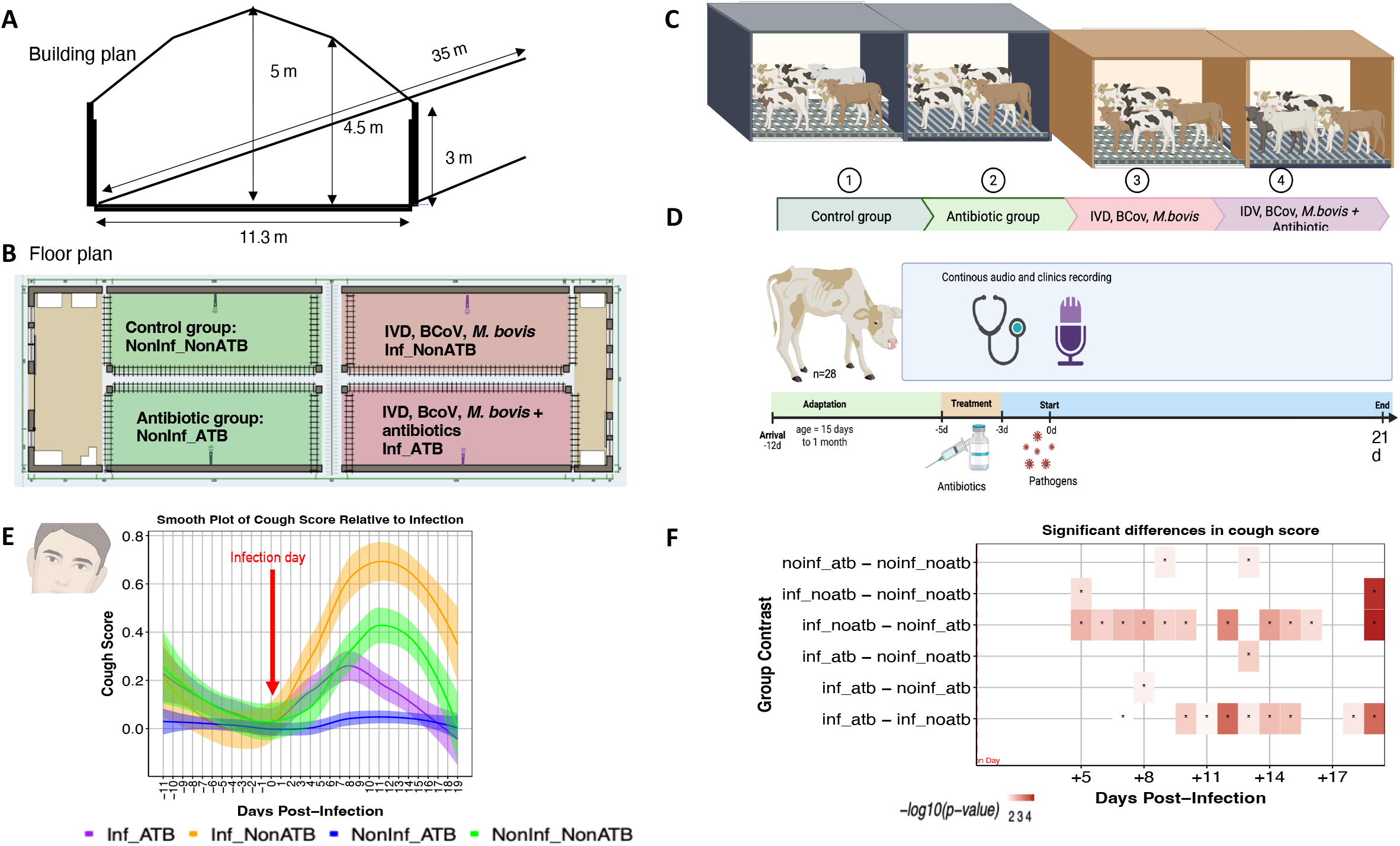
Performance and deployment metrics of the acoustic monitoring system. (A) Total hours of audio recorded by each sensor across the 31-day experimental period. Bars represent cumulative recording time per sensor, aligned to the experimental timeline; (B) Training dynamics of the binary classifier. Top: training loss per epoch; Bottom: training accuracy and F1-score per epoch. Both plots span 75 epochs (x-axis: 0–70 shown for clarity), illustrating rapid convergence and stable performance; (C) Confusion matrix displaying the performance of the sound classification model for detecting cough events from microphone recordings. Rows represent true labels, and columns indicate predicted classes. Values indicate the number of true and predicted labels for cough and non-cough events; (D) Mean posterior probability of cough classification across all processed files, stratified by sensor. Lines represent the average model confidence for positive predictions, reflecting consistency across deployment sites.

This extensive deployment yielded a large, acoustically diverse dataset that served as the basis for supervised model training and performance evaluation. After preprocessing and quality control, we developed a binary classifier using 1,045 manually annotated audio clips (308 cough; 737 non-cough). Training over 75 epochs (batch size□=□16; learning rate□=□0.001) resulted in rapid convergence, with loss decreasing from approximately 32.5 to 15 and the F1-score stabilizing near 90% (Fig.□2B). The final model demonstrated strong performance: overall accuracy reached 0.92, with macro and weighted F1-scores of 0.91 and 0.92, respectively. For non-cough events, precision, recall, and F1-score were 0.94, 0.95, and 0.95, while for coughs all three metrics reached 0.87, indicating high sensitivity despite substantial acoustic variability and an imbalanced class distribution (Fig.□2C). A posterior-probability threshold of 0.85 was selected for cough classification (Fig.□2D), providing a biologically meaningful and conservative cutoff for event detection.

To mitigate class imbalance, we incorporated a class-weighted loss function during training. The strongly weighted performance metrics further suggest that the inherent acoustic distinctiveness of cough events helped counterbalance the unequal class representation.

Building on this validated model, we applied the classifier to the full dataset, comprising 35,786 processed audio segments. Of these, 6,868 (19.2%) were classified as coughs, corresponding to an average of 221.5 coughs per day and 2.5 coughs per hour, while 28,918 (80.8%) were classified as non-coughs. These results confirm the classifier’s ability to robustly distinguish coughs from background acoustic signals, including frequent metallic noises generated by gates, fences, and feeding equipment within the calf rearing environment.

### Acoustic Cough Surveillance between pens: Spatial Accuracy and Biological Sensitivity

To evaluate group-level differences and temporal dynamics in acoustic signal reliability, cough activity across pens and time points was analysed. The classifier’s overall performance was first assessed across the groups. Event frequency varied considerably across groups, whereas detection performance was consistent across sensors, with each reporting a mean confidence above 90% (Table 1).

**Table 1.**
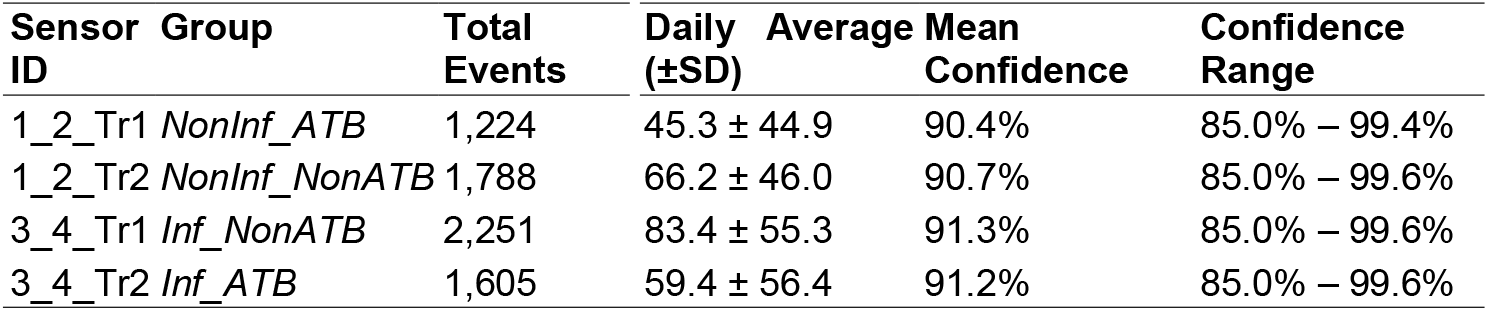
Sensor-level analysis of the HuBERT cough-detection model.

Sensor *Inf_NonATB* recorded the highest proportion of events (32.8%), while *NonInf_ATB* reported the fewest events (17.8%; Table 1). To further assess the relationship between pen-level infection status and recorded cough frequency, a Poisson regression model was fitted using event count as the outcome variable. Using the *NonInf_NonATB* as the reference category, all groups demonstrated statistically significant differences in cough event frequency. The *Inf_NonATB* exhibited the highest activity, with 67% more detected events (β□=□0.51; exp^(β)□^=□1.67). Conversely, *NonInf_ATB* sensors registered 44% fewer events compared to the reference (β□=□– 0.59; exp^(β)^□=□0.56). A significant temporal trend was also observed, with each additional day from day 11 associated with a 3.5% reduction in event detection (β□=□– 0.036; exp^(β)^□=□0.965), reflecting recovery over time. All effects were highly significant (*p*□<□0.001).

Temporal analysis revealed notable day-to-day variation (SD = 141.7 events/day), with the highest cough activity observed on 9 days post-infection (522 events) and the lowest on the last day of the experiment (30 events; Fig. 3A). Particularly, for experimentally infected groups, the highest counts were recorded by sensor *Inf_NonATB* on Day 9 post-infection (247 events) and sensor *Inf_ATB* on Day 8 post-infection (189 events; Fig 3AB). Non-infected groups also demonstrated periodic spikes, notably on 13-14 days post-infection (226 and 208 events, respectively). During this process, a technical anomaly was identified on day 12 post-infection (Fig. 3B). On this date, one calf in the *NonInf_NonATB* group remained inadvertently attached to the mobile feeding fence used for milk distribution, producing continuous mechanical and metallic sounds that dominated the acoustic recordings. These non-biological signals likely interfered with the classifier’s ability to accurately detect cough events (Fig. 3B).

**Figure 3.**
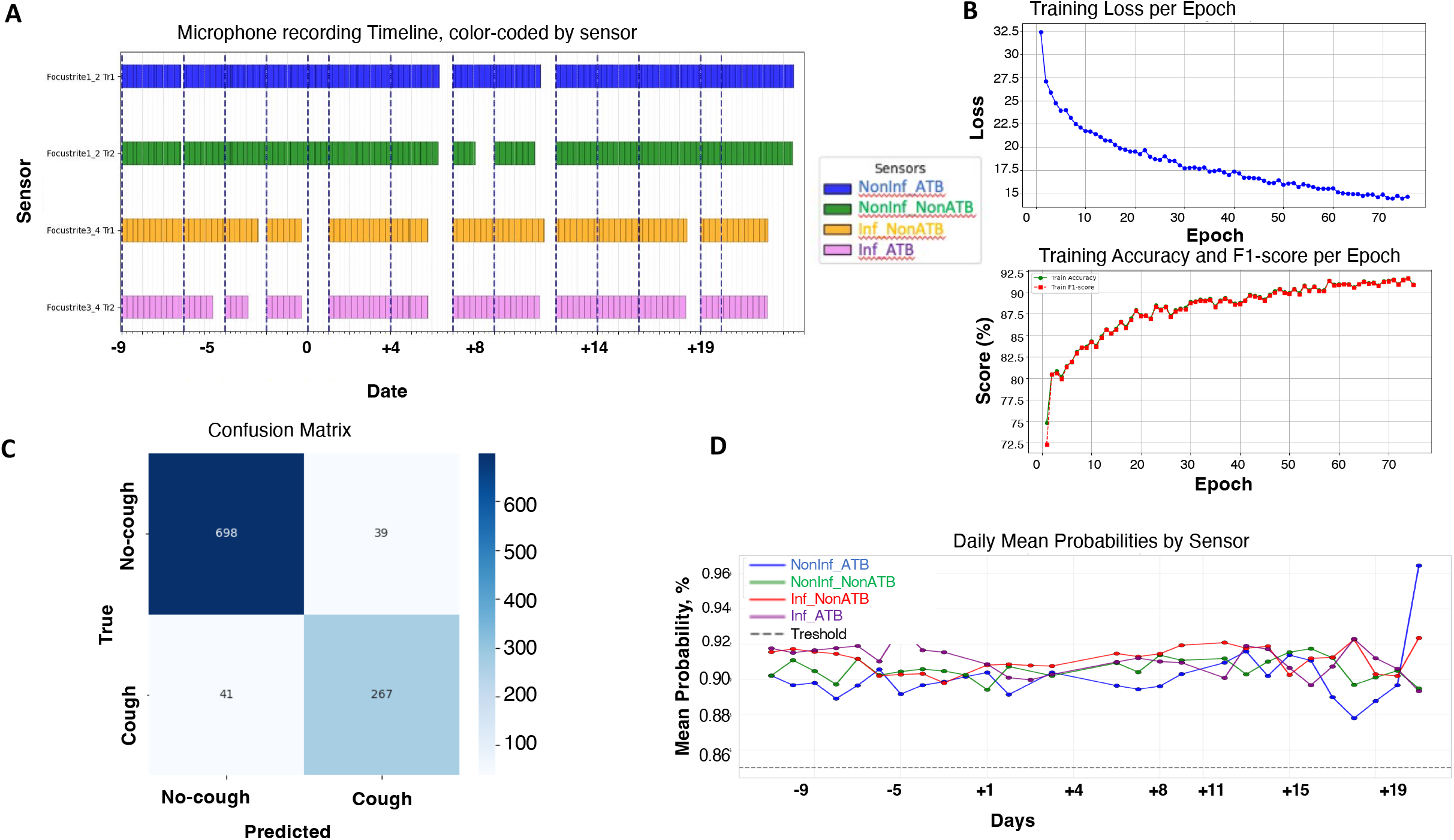
Cough Score and Audio Event Frequency Aligned by Days Post-Infection. (A) Event Count Heatmap depicting daily detection frequencies across four sensor channels using a classification threshold of ≥0.85. Darker shades indicate higher event counts. This visual representation allows for temporal and spatial comparison of sound event detection; (B) Temporal patterns of audio event frequency, aligned by days post-infection across four experimental groups; (C) Circadian pattern analysis of cough detection events with kernel density smoothing. Individual events are plotted as scatter points along the 24-hour timeline (*x*-axis) for each sensor (*y*-axis), with random vertical jitter to prevent overlap. Coloured trend lines show kernel density estimation of activity patterns, with filled areas representing activity intensity; (D) Time-series plots display daily cough event counts for each sensor group, with the *x*-axis representing experimental days relative to infection (Day 0) and the *y*-axis showing event frequency. Lines indicate daily event trajectories. Red dashed lines denote group-specific alarm thresholds, calculated as *μ* + 2*σ*, where *μ* is the mean baseline cough count (Days –5 to 0) and *σ* is the corresponding standard deviation. Red points highlight alarm-positive days where event counts exceeded the threshold, and “Alarm” labels annotate these events. All panels share a fixed *y*-axis scale to facilitate cross-group comparison.

In addition to infection-related dynamics, acoustic monitoring also revealed a noticeable rise in respiratory sound activity during the initial days of the experiment. As mentioned above, this early peak in cough events likely stemmed from management-related stressors, particularly the regrouping of unfamiliar animals and the exposure to fresh pine shavings. Thus, despite the close physical proximity between pens, the acoustic monitoring system consistently captured pen-specific respiratory activity with high confidence, revealing clear spatial discrimination between sensors and significant variations in cough frequency driven by infection status, treatment, time, and management patterns.

Interestingly, event detection peaked during early morning and late evening hours for most sensors (Fig. 3C). These event trends, visualized with smoothed lines and shaded confidence intervals, likely echoed patterns of animal behaviour. Overall, minimal event activity was observed between 00:00 and 03:00, aligning with expected resting phases.

### Acoustic Monitoring as an Early-Warning Tool: Evidence for Predictive Cough Detection Prior to Clinical Onset

To assess the alignment between AI-based acoustic monitoring and veterinarian cough scores detection, we applied generalized additive models across all treatment groups. The temporal trajectory of cough scores assigned by veterinarians did not significantly correlate with the acoustic signal patterns in any group, including *NonInf_NonATB* (edf = 1, F = 0.011, *p* = 0.919), *Inf_NonATB* (edf = 1.72, F = 0.915, *p* = 0.392), *NonInf_ATB* (edf = 1.47, F = 0.411, *p* = 0.675), and Inf_ATB (edf = 1.00, F = 0.002, *p* = 0.963). Notably, in the *Inf_NonATB* group, microphone-detected cough activity began rising 1–2 days prior to cough score escalation, indicating that acoustic signals may precede observable clinical signs.

To further investigate the anticipatory capacity of acoustic monitoring, an alarm thresholding system was implemented based on pre-infection cough data. For all sensor groups, baseline activity was established using event counts recorded from Days -5 to 0 relative to the infection date. The resulting average threshold was 112 coughs/day. Days exceeding this value were flagged as alarm-positive, indicating anomalous increases in cough activity. Post-infection alarm activation occurred in the *Inf_ATB* group on Days 7 and 8, and in the *Inf_NonATB* group on Days 7, 8, 11, and 12. Notably, alarm-positive days were also detected in the non-infected groups on Days 12 and 13. These findings support the potential of acoustic surveillance to detect early behavioural indicators of respiratory compromise prior to the onset of clinically scorable symptoms (Fig. 3D).

As previously noted, the alarm threshold was exceeded during the initial days of the experiment, particularly on Day –9, across multiple groups. These early activations occurred prior to infection and baseline calibration, and all animals tested negative for major respiratory pathogens upon arrival. This suggests that the elevated cough activity was likely non-infectious and instead associated with management-related factors.

## Discussion

Early detection of respiratory disorders in calves is critical for managing BRDC, yet conventional clinical scoring methods often detect illness only after substantial physiological compromise has already occurred. In this study, we demonstrate that microphone-based, AI-powered acoustic monitoring provides a sensitive, automated, real-time, and non-invasive window into respiratory health, capturing cough dynamics with a temporal resolution far exceeding that achievable through manual observation. By continuously monitoring calves experimentally co-infected with Influenza D virus, Bovine Coronavirus, and *Mycoplasma bovis*, the system revealed a clear and biologically coherent trajectory of cough expression—emerging around Day +5, peaking near Day +11, and declining thereafter. This pattern aligns with the established pathogenesis of BRDC, in which early viral infection primes the respiratory tract for subsequent bacterial colonisation and clinical exacerbation (Cummings et al., 2022; Gaudino, Chiapponi, et al., 2022; Griffin et al., 2010b; Salem et al., 2019b).

Importantly, the system also captured the therapeutic impact of pre-infection tilmicosin, with treated calves showing a marked reduction in cough frequency from Day 9 onward. This dual capability positions microphone-based monitoring not only as an early-warning tool but also as a valuable instrument for evaluating treatment efficacy and supporting evidence-based antimicrobial stewardship.

Spatial analysis further validated the acoustic system’s robustness. Each microphone reliably captured pen-specific coughs with minimal cross-detection and classification confidence exceeding 90%, demonstrating strong localisation capacity despite the inherently noisy barn environment. This precision is essential for commercial settings, where multiple pens, high stocking densities, and overlapping sound sources can complicate signal attribution. Notably, the system anticipated the rise in clinical symptomatology by approximately two days—an early-warning advantage consistent with findings from other Precision Livestock Farming (PLF) technologies such as accelerometers and smart feeders, which detect behavioural deviations prior to overt clinical signs (Bushby et al., 2024; Kim et al., 2024). The integration of a cough-frequency-based alarm system operationalised this early-warning potential, with alarms triggered up to 2 days before peak clinical scores, particularly in the infected and untreated group. Such anticipatory signals are valuable for enabling timely, well-informed management decisions that can reduce disease severity, improve clinical outcomes, and ultimately promote calf health and welfare.

From a technical perspective, our approach builds on and extends previous work in bovine acoustic monitoring (Carpentier et al., 2018; Kim et al., 2024; Vandermeulen et al., 2016). Earlier studies demonstrated promising performance for calf-cough detection using handcrafted acoustic features or complex model architectures, typically achieving precision and recall in the 0.80–0.90 range. Our HuBERT-based pipeline achieved comparable or superior performance while providing a more streamlined, scalable framework. HuBERT, originally developed for human speech representation learning, captures rich hierarchical acoustic features (Hsu et al., 2021). Its ability to encode subtle temporal and spectral patterns proved advantageous for distinguishing coughs from diverse barn noises, even under challenging acoustic conditions. The use of short, non-overlapping 1-second windows enabled fine-grained temporal tracking, while a conservative decision threshold (≥ 0.85) and notch-based noise augmentation enhanced robustness during deployment.

## Conclusion

Overall, our findings demonstrate that microphone-based acoustic monitoring is a powerful and practical tool for early detection of respiratory disease in calves, offering continuous, objective, and high-resolution surveillance that complements and enhances traditional clinical assessment. The strong performance of the HuBERT-based classifier, combined with the system’s ability to anticipate clinical symptomology and quantify treatment response, underscores the potential of acoustic PLF technologies to enhance respiratory health management. As farms continue to scale and labour availability declines, such automated monitoring systems will become increasingly indispensable for safeguarding animal health and welfare, reducing antimicrobial use, and improving the sustainability of livestock systems.

## Limitations of the study

Despite the promising results, our study has several limitations that warrant consideration. First, the sample size and pathogen diversity were limited, restricting the generalizability of our findings across different herds, breeds, and epidemiological contexts. Additionally, the housing configuration and microphone placement were optimized for this setup; acoustic propagation and signal discrimination may vary in larger or differently designed facilities. Additionally, it cannot detect respiratory problems in individual animals.

While HuBERT has shown strong performance in human acoustic tasks, its application in livestock remains novel, and further benchmarking against alternative self-supervised models is needed. Future studies should incorporate air quality metrics and behavioural observations to better disentangle pathological and environmental signals. Moreover, beyond mere detection, cough sounds contain distinct latent features—such as duration, peak frequency [Hz], and spectral shape—that can be exploited to classify cough types and associated vocalizations (Ijaz et al., 2022). These acoustic traits may provide deeper insight into health and welfare status, potentially distinguishing physiological coughs linked to disease severity, pain, or distress. In this way, characterizing cough types could also support more informed decision-making regarding whether antimicrobial treatment is warranted or whether alternative management strategies are more appropriate. To fully harness this diagnostic potential, future AI-acoustic models should evolve beyond binary classification and incorporate richer acoustic profiling.

## Ethics approval

The local animal care and use committee and the French Minister of Research reviewed and approved the study protocol for the calves’ study (reference: APAFIS #49942-2024062000479999 v3, dated September 23rd, 2024, Ethics Committee no. 115). All protocols were conducted in accordance with EEC Regulation (No. 2010/63/UE) governing the care and use of laboratory animals, which has been effective in France since January 1, 2013.

## Data and model availability statement

Examples of raw acoustic recordings, pre-processed datasets, and the model checkpoint used in this study are available via the INRAE data repository (https://entrepot.recherche.data.gouv.fr/dataverse/inrae), under the DOI [https://doi.org/10.57745/FP5LA9].

The repository includes four raw audio files, two from each of two nights, recorded at the ENVT experimental facility using directional microphones. Each file spans approximately 6-7 hours and reflects the unprocessed acoustic environment of the pens during nighttime recording sessions. In addition, the dataset also includes a curated subset of 40 short audio clips: 20 manually annotated cough events and 20 non-cough events, representing a range of cough events and barn sounds. These clips were used in the training and evaluation of the detection algorithms. The repository also contains a HuBERT-based classifier checkpoint (.pth) saved during early training iterations (no training or inference implementation is included). All materials are provided for research and educational purposes only. Users are encouraged to cite the corresponding publication and acknowledge the authors when using these resources.

For access to the full event-level dataset or further technical details, interested researchers may contact Ignasi Nou-Plana at.

## Declaration of generative AI and AI-assisted technologies in the writing process

Generative AI tools, including Microsoft Copilot, were used to assist with editing and refining the manuscript. The authors reviewed and approved all content to ensure accuracy and scientific integrity.

## Acknowledgement

We sincerely thank the PhD students, postdoctoral researchers, technicians, and veterinary students who ensured the daily care and welfare of the animals throughout the experiment. In particular, we are grateful to Maverick Monie--Ibanes, Thomas Peyret, Guenaelle Derolez, Fatima-Zohra Sikht, Corto Martin-Rebouissou, Romain Bonsergent, Valentin Puech, David Bars, Joshua Malsa, Gwendoline Pot, Hortensia Robert, and Florence Mompart for their dedication and support during the study.

---

**Figure S1.**
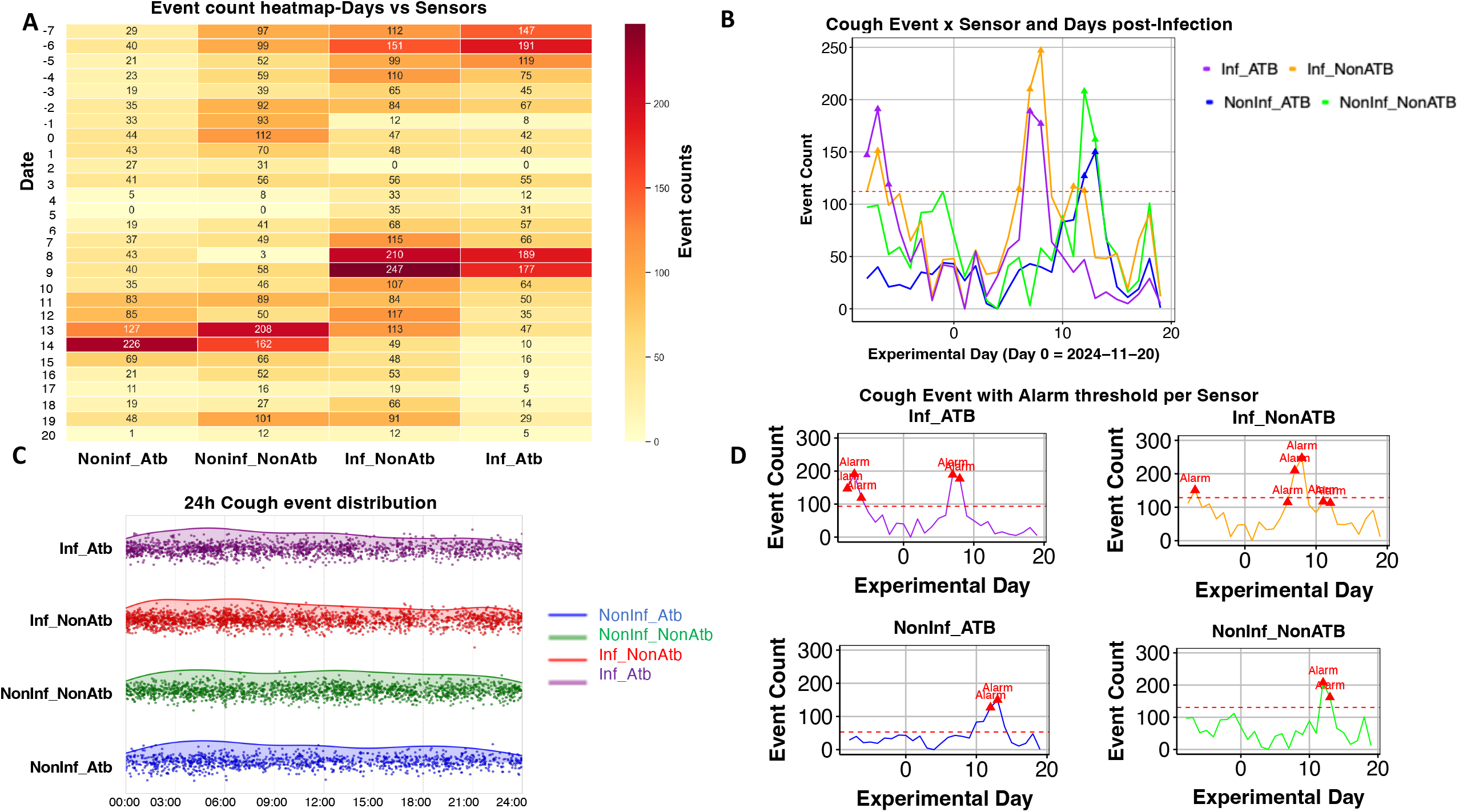
Environmental Stability and Feeding Behaviour Across Experimental Groups. (A) Environmental conditions between calves in control pens (green lines) and infected pens (purple lines), including dew point, heat index, relative humidity, temperature-humidity indices, and ambient temperature. Shaded zones indicate thresholds for thermal comfort and stress, such as “Caution,” “Dry & Crisp,” “Extreme Caution,” and “Tropical & Oppressive.” These parameters remained relatively stable across pens and time, suggesting that environmental fluctuations did not confound respiratory outcomes; (B) Milk consumption and refusal, with separate lines for four experimental groups. A vertical dashed red line marks the day of infection. This panel illustrates the dynamics of feeding behaviour in relation to infection status and treatment, providing insight into clinical progression and recovery.

