## Supplementary figures and images for "AI-Powered Acoustic Surveillance for Early Detection of Calf Respiratory Disease"

### Figure S1

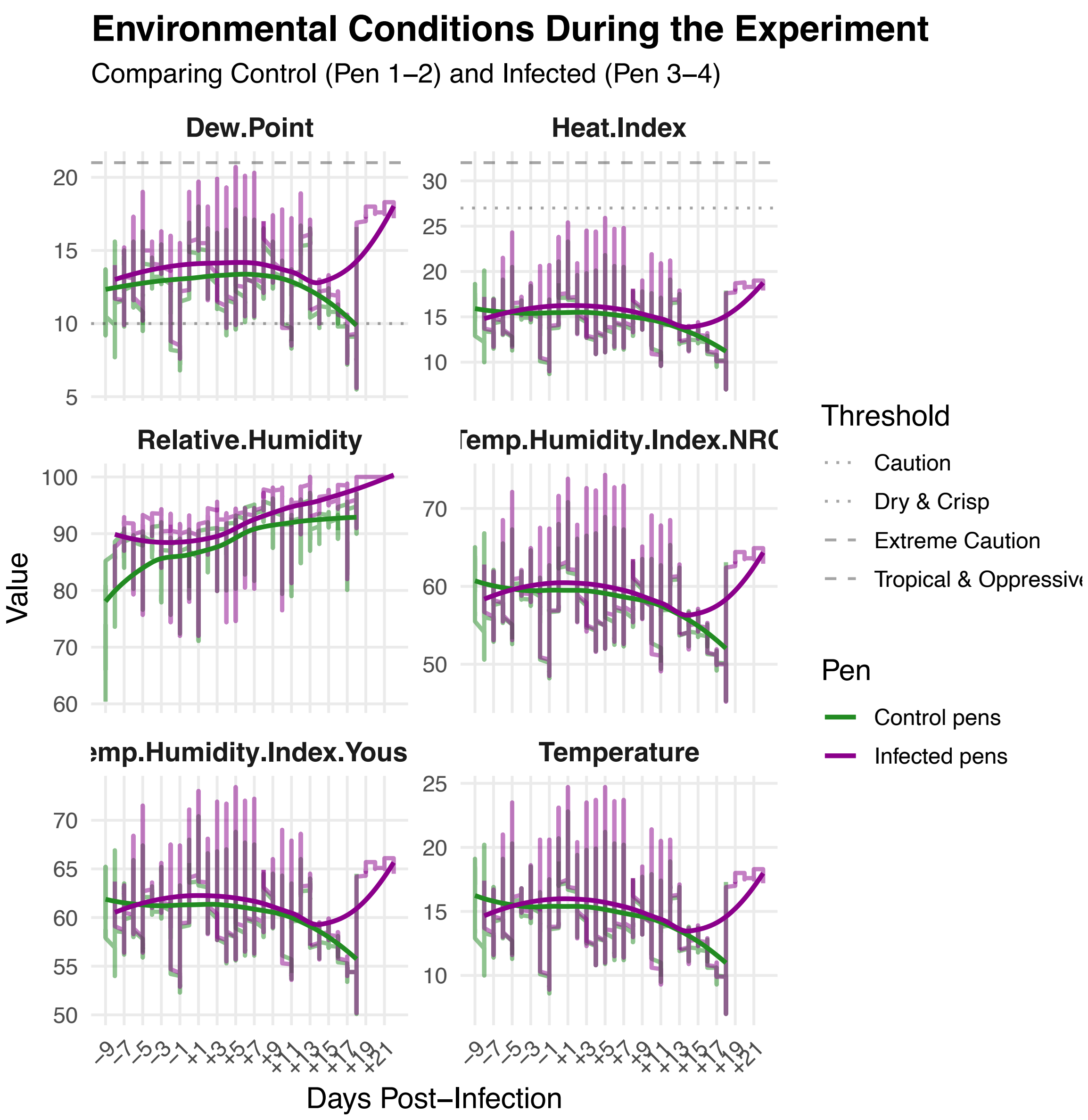

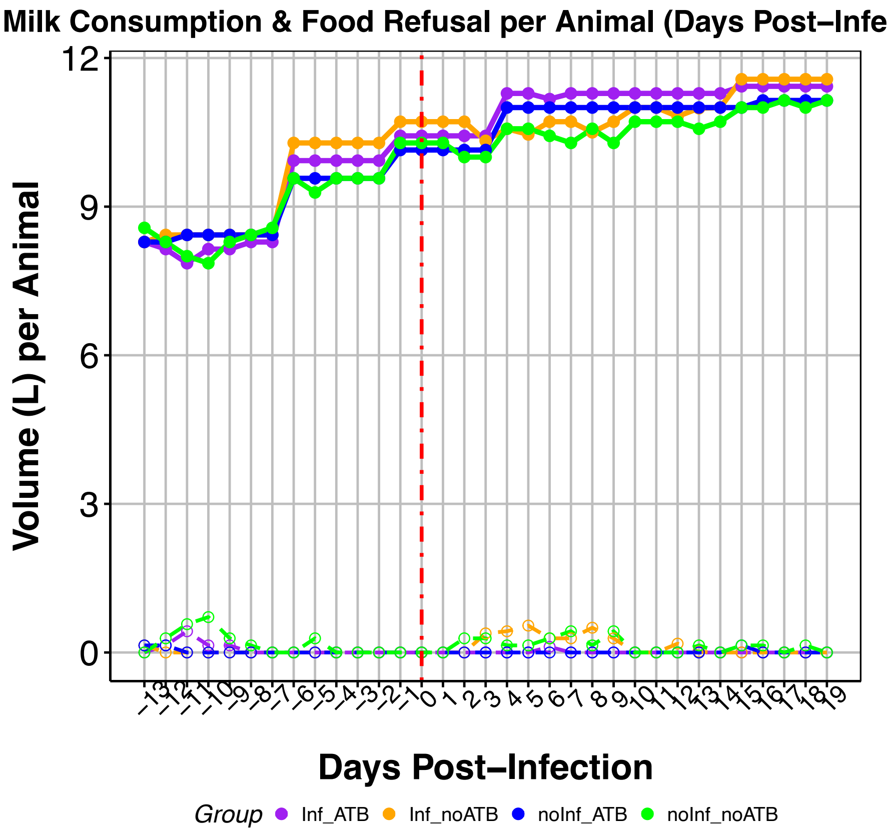
